# An accelerated molecular dynamics study for investigating protein pathways using the bond-boost hyperdynamics method

**DOI:** 10.1101/2024.09.30.615751

**Authors:** Soon Woo Park, Moon-ki Choi, Byung Ho Lee, Sangjae Seo, Woo Kyun Kim, Moon Ki Kim

## Abstract

Molecular dynamics (MD) simulation is an important tool for understanding protein dynamics and the thermodynamic properties of proteins. However, due to the high computational cost of MD simulations, it is still challenging to explore a wide conformational space. To solve this problem, a variety of accelerated MD schemes have been proposed over the past few decades. The bond-boost method (BBM) is one of such accelerated MD schemes, which expedites escape events from energy basins by adding a bias potential based on changes in bond length. In this paper, we present a new methodology based on the BBM for accelerating the conformational transition of proteins. In our modified BBM, the bias potential is constructed using the dihedral angle and hydrogen bond, which are more suitable variables to monitor the conformational change in proteins. Additionally, we have developed an efficient algorithm compatible with the LAMMPS package. The method is validated with the conformational change of Adenylate kinase (AdK) by comparing the conventional and accelerated MD simulation results. Based on the accelerated MD results, the characteristics of AdK are investigated by monitoring the conformational transition pathways and the behavior of interdomain salt bridges. Moreover, the free energy landscape calculated using umbrella sampling confirms all the states identified by the accelerated MD simulation are the free energy minima and the system makes transitions following the path indicated by the free energy landscape. Our efficient approach is expected to play a key role in investigating transition pathways in a wide range of protein simulations.

## Introduction

Proteins perform their biological functions through conformational changes such as side-chain isomerization, domain motions, and allosteric regulations. As proteins play various roles in vivo (e.g. transport of metabolites, chemical concentration regulation, and catalysis), investigating their conformational changes is essential to comprehend their underlying biological phenomena. Over the past few decades, numerous experimental and computational studies have been performed to elucidate these changes, among which computational investigations have been mainly conducted with molecular dynamics (MD) simulations [1, 2, 3, 4]. MD simulation is one of the most widely used simulation tools because it provides a time-dependent evolution and samples the conformational space of a system by numerically calculating Newton’s equation of motion. However, despite the advances in computer power, conducting simulations of fully-atomistic models with realistic interatomic potentials remains computationally expensive, leading to limited simulation times on the nano- and micro-second timescales. Additionally, biological phenomena of interest, such as protein folding, large conformational change, and protein-ligand interaction, take place as rare events over a long timescale because biomolecular systems are trapped in their local energy minima most of the time during the simulation. To address this issue, a wide range of enhanced sampling techniques have been developed, including umbrella sampling [5], metadynamics [6], adaptive biasing force [7, 8, 9], steered MD [10], and replica-exchange [11].

One such accelerated technique is hyperdynamics, which extends the timescale of an MD simulation without any advanced knowledge of the system such as the dividing surfaces or the neighboring states [12, 13]. In hyperdynamics, the escape event from an energy basin is accelerated by adding a predefined bias potential to the system. To accomplish this, the bias potential is constructed in the region excluding the transition state by calculating the eigenvalues of the Hessian matrix of the system. However, because the eigenproblem of the Hessian matrix, which is computationally expensive, needs to be solved at every step, the computational overhead dramatically increases as the system size or simulation time increases. To address this problem, several variants of hyperdynamics have been developed, utilizing less computationally intensive variables to define bias potentials [14, 15, 16, 17, 18]. Among them, the bond-boost method (BBM) is a variant of the hyperdynamics method in which the change in bond length is used to construct the bias potential [14]. In the BBM, the status of the system (i.e. in-state or out-of-state) is determined solely based on a predefined criteria related to the bond length without calculating the eigenvalue of the Hessian, thus resulting in much less computational overhead than the original hyperdynamics method for long-time simulations. While the BBM has been successfully applied to a number of material systems [19, 20, 21, 22], the structural complexity of proteins has still hampered the applicability of the BBM to biomolecular systems.

Here, we introduce a modified version of the BBM that is optimized for investigating the conformational pathway of proteins. The main difference of our method from the original BBM is that we use the dihedral angles affecting the conformational transition and the hydrogen bonds connected to a ligand as the boost targets. We have also developed an efficient algorithm for accelerating conventional MD simulations by optimizing several key parameters, such as maximum bias potential and threshold, based on the characteristics of the proteins.

To validate our method, we chose Adenylate kinase (AdK), a monomeric phosphotransferase enzyme that has attracted much attention as an ideal testbed for the application of a new pathway sampling method [23, 24, 25, 26, 27]. It is widely known that AdK undergoes a large conformational change of domains during the catalytic cycle, which regulates cellular energy homeostasis within the cell by catalyzing the reaction ATP MG^2+^ +AMP ↔ ADP MG^2+^ +ADP. Since alterations in AdK’s activity can lead to diseases such as hemolytic anemia, metallic disorders, and cancer, it is crucial to identify AdK’s conformational pathway related to the catalytic cycle.

Currently, various structures of AdK have been revealed mostly by experimental studies. Among them, the apo (substrate-free) form (PDB ID: 4AKE) [28] and substrate binding form (PDB ID: 1AKE) [29] have been popularly used in computational studies as the open and closed representative structures, respectively. The structure of AdK is composed of three domains: an ATP-binding (LID) domain (residues 122–159), a CORE domain (residues 1–29, 60–121, and 160–214), and an AMP-binding (NMP) domain (residues 30–59) (Figure 1). Among them, the LID and NMP domains are dynamic structures relative to the CORE domain and play a key role in binding substrates through a large conformational change over a long timescale. Therefore, in this study we investigated the transition of AdK from the closed structure to the open structure with our accelerated MD technique from the perspective of the conformational pathway for domain movements. We performed both conventional and accelerated MD simulations for comparison and found that only the accelerated MD simulations exhibited transitions from the closed structure to the open structure, exploring a broader conformational space than conventional MD simulations.

**Figure 1.**
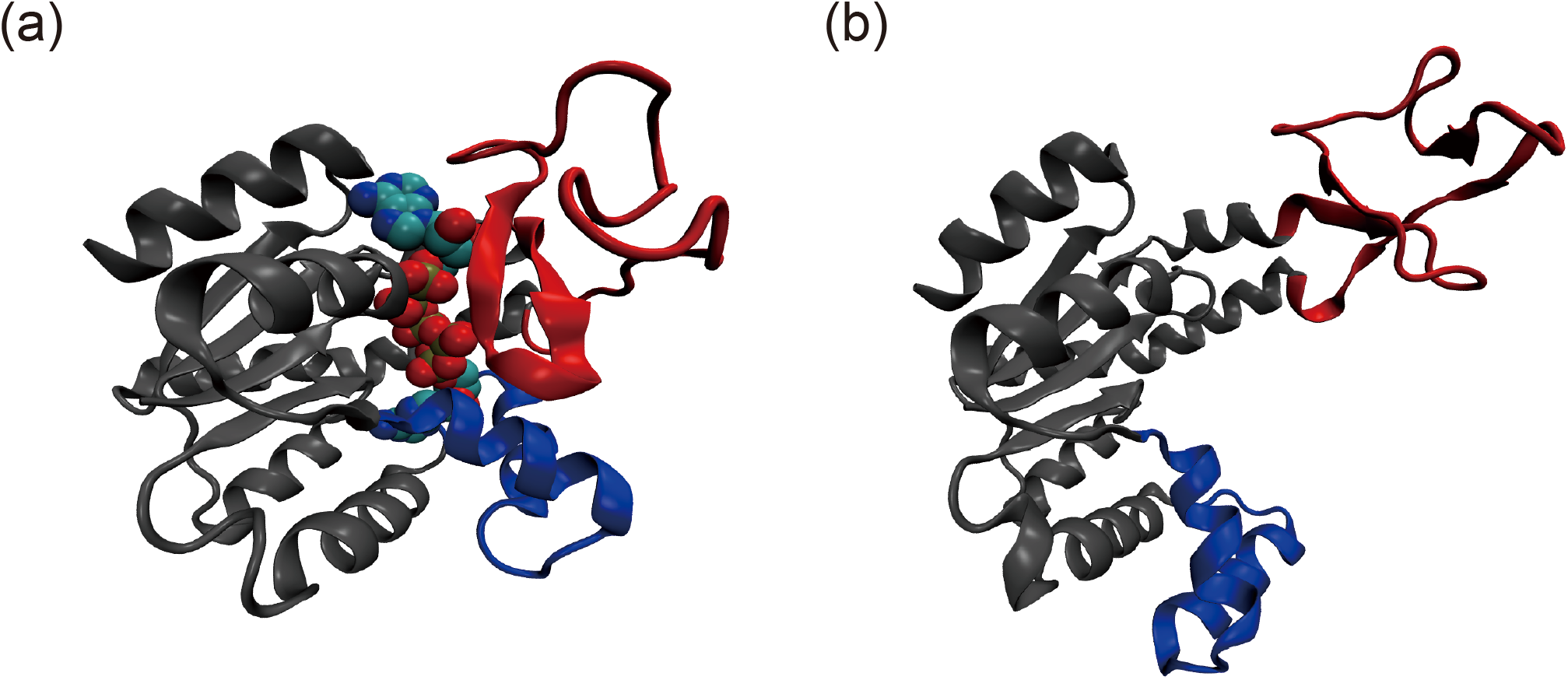
The structure of Adenylate kinase can be described by two distinct conformations, namely the (a) closed (PDB ID: 1AKE) and (b) open (PDB ID: 4AKE) forms. In the closed structure, an inhibitor called Ap5A, represented using the van der Waals (vdW) spheres, is visibly bound to the interior of the protein. The protein consists of three domains: an ATP-binding (LID) domain (red), a CORE domain (gray), and an AMP-binding (NMP) domain (blue). These two conformations have been visualized and rendered using the VMD program [30].

## Methods

### Bond-boost method (BBM)

In the BBM, an empirical bias potential Δ*V* is defined as

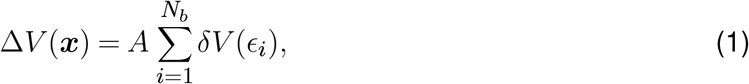

where *x* collectively represents the positions of all the tagged atoms, *A* is the envelope function, *N*_*b*_ is the number of tagged bonds, and *ϵ*_*i*_ is the fractional change in the bond length of the tagged bond *i*, and *δV* is the local boost potential determined by *ϵ*_*i*_. All the atoms related to the transition are tagged and all the nearest-neighbor bonds of the tagged atoms are boosted by the local boost potential. *ϵ*_*i*_ is defined as

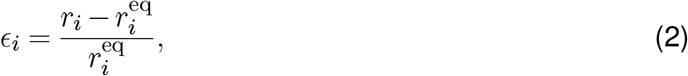

where *r*_*i*_ and 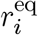 are the current and equilibrium bond lengths of the tagged bond *i*, respectively.

In hyperdynamics methods, such as the BBM, it is critical that the original potential is not affected by adding the bias potential near the dividing surface to prevent any distortion of the natural pathways. Assuming that *ϵ*_*i*_ will increase as the system approaches the dividing surface and the transition state, a threshold value *q* is introduced so that *δV* vanishes when |*ϵ*_*i*_| ≥ *q*. The following function form was used in the original BBM.

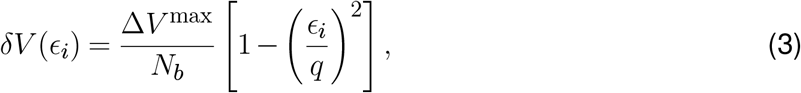

where Δ*V*^max^ is the maximum boost energy. If the threshold *q* is too small, the boost effect is insignificant because the range of the boost is too narrow. Conversely, if the threshold *q* is too large, the boost potential may affect the dividing surface and the transition states so that it can distort the natural pathway of the system. Also, the maximum boost energy Δ*V*^max^ should be chosen as a value that does not exceed the natural energy barrier of the system. A larger Δ*V*^max^ gives a greater boost effect, but adding too much boost to the local minimum may create a new local maximum near the equilibrium position, causing the bias potential to destroy the natural shape of the potential energy surface.

Even though *δV* becomes zero whenever its corresponding bond length change exceeds the threshold value, the total bias potential Δ*V* may not be zero unless all the tagged bonds exceed the threshold simultaneously. To remedy this problem, the envelope function, which becomes zero if any of the *ϵ*_*i*_ values is greater than *q*, is multiplied to the sum of the local boost potentials as seen in Eq. (1). This is achieved by defining *A* as a function of *ϵ*^max^ = max {|*ϵ*_*i*_|}:

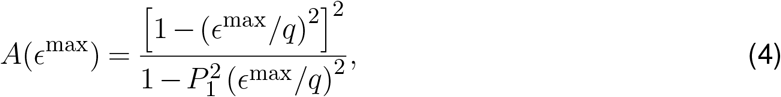

where *P*_1_ is a parameter introduced to make the derivative of the function continuous at the boundary of *ϵ*^max^ = *q*, typically ranging from 0.9 to 0.98.

The use of the envelope function has a disadvantage of reducing the overall boost effect of the system. Specifically, the envelope function decreases to zero even when only one tagged bond exceeds the threshold value while all the other bond lengths are still far below the threshold. Especially, in large systems, not boosting the entire system due to an accidental cross-over of a single bond can significantly reduce the efficiency of the BBM. In an effort to overcome this problem, a local hyperdynamics method was proposed [34], where the boosted region is separated into multiple sub-regions, assuming little correlation between them, and a local bis potential is applied to each sub-region independently from each other. Here, we adopted the idea of the local hyperdynamics method to effectively apply the BBM to biosystems.

### Modification of the bond-boost method

To properly accelerate the dynamic evolution of a biomolecular system, several parameters that comprise the bias potential need to be redefined, including the boost target, equilibrium position, threshold, and maximum boost energy. First, we have chosen as the boost targets the dihedral angle and the hydrogen bond to adequately monitor conformational changes in proteins. Compared to bond lengths and bond angles, dihedral angles are more flexible and exhibit larger changes during the transition and thus were chosen as the boost target in our method. Moreover, we selected only those dihedral angles forming the backbone positioned at hinge regions because the dihedral angles constituting secondary structures and domain regions remain rigid during the conformational transitions. Additionally, hydrogen bonds between the protein and ligand were chosen as the boost targets considering that phenomena occurring between the protein and ligand can be effectively accelerated by weakening the bonds between them. Our approach can be easily extended even for large proteins by effectively adding the boost targets that are characteristic to those systems. This is a significant advantage when predicting protein conformational pathways.

The quadratic function employed for the local boost potential in the original BBM (Eq. (3)) creates a discontinuous force at the boundary where the energy becomes zero. To prevent abrupt alterations in the force at the boundary, we adopted the functional form similar to the envelope function form including the smoothing parameter (Eq. (4)). The corresponding equations are

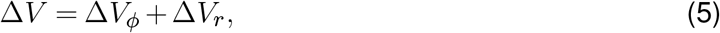

where

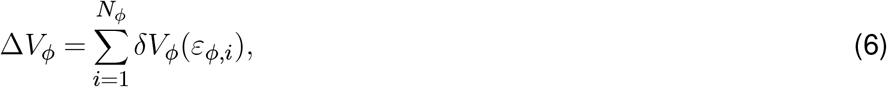

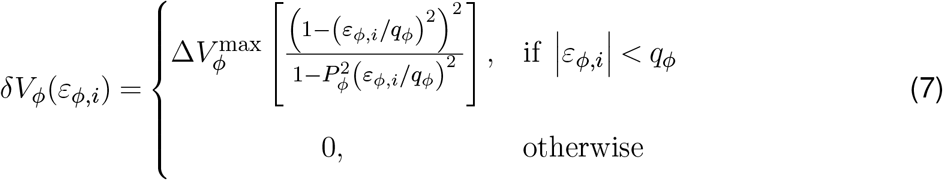

where *N*_*ϕ*_ is the number of targeted dihedral angles, 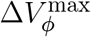 is the maximum boost energy for dihedral angle, *P*_*ϕ*_ is the parameter that ensures a zero gradient at the boundary, *q*_*ϕ*_ is the threshold for dihedral angle, and 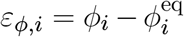 with 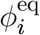 being the equilibrium dihedral angle, and

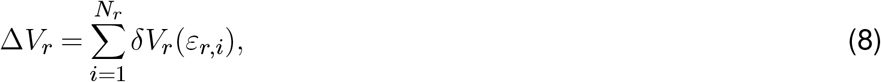

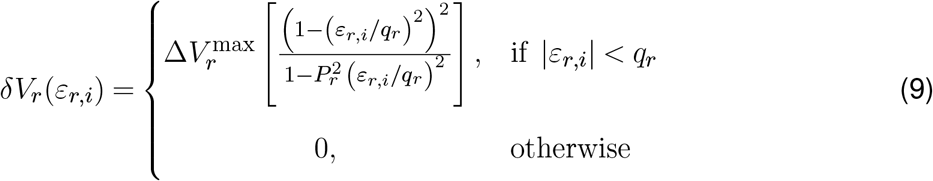

where *N*_*r*_ is the number of targeted hydrogen bonds, 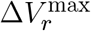 is the maximum boost energy for hydrogen bond, *P*_*r*_ is the parameter that ensures a zero gradient at the boundary, *q*_*r*_ is the threshold for hydrogen bond, and 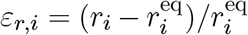 with 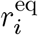 being the equilibrium hydrogen bond length.

As seen in Eqs. (6) and (8), we adopted the local hyperdynamics method which boosts each individual boost target separately. To ensure the validity of this idea, we checked the correlation between them over time and found that all the boost targets are sufficiently uncorrelated. Moreover, as will be seen in the results and discussion section, the transition pathway from the accelerated MD using our method followed the natural pathway of the system on the free energy surface.

Next, to determine the equilibrium position of each boost target, which characterizes the local minimum of the system, we introduced a new concept called the waiting time *t*^waiting^. In the BBM, the equilibrium position of the target is reset by performing a conjugate-gradient minimization every specific time steps while no boosting occurs (i.e. the system is out of the state). While this approach is computationally efficient for some systems, it may not be practical for larger and more complex systems due to the increased computational demand. Additionally, the minimization technique does not consider the effect of temperature. In our approach, we monitor a boost target (a dihedral angle or a hydrogen bond length) for the time period *t*^waiting^ during an MD simulation at a given temperature and calculate the average value regarded as the equilibrium position (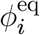or 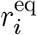), which provides a more accurate representation of the system’s behavior at the given temperature. To ensure that the equilibrium position sufficiently converges, we set *t*^waiting^ to be 100 ps for the dihedral angle and the hydrogen bond in our method.

Finally, the threshold (*q*_*ϕ*_ and *q*_*r*_), which is the criterion for determining the transition, and the maximum boost energy (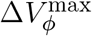 and 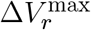), which determines the magnitude of boost, were deduced by examining the probability distribution of each boost target. Figures 2a and 2b illustrate the representative examples of the behaviors of the boost targets during a transition and their resultant probability density distributions, respectively. The threshold should be set to clearly distinguish the states before and after the transition. After examining all the probability distributions, *q*_*ϕ*_ and *q*_*r*_ were set to 40° and 0.2, respectively. Here, the thresholds were empirically chosen as values that do not alter the natural transition pathway. Moreover, we determined 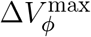 and 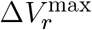 by considering the change in the probability density distribution due to the application of the bias potential, using the method developed by Kim and Falk [18]. Our criterion is not to introduce additional barriers inside the state which may lead to a different evolution from the original system. The chosen values for 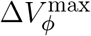 and 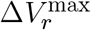 are 1 kcal/mol for the dihedral angle and 3 kcal/mol for the hydrogen bond, respectively.

**Figure 2.**
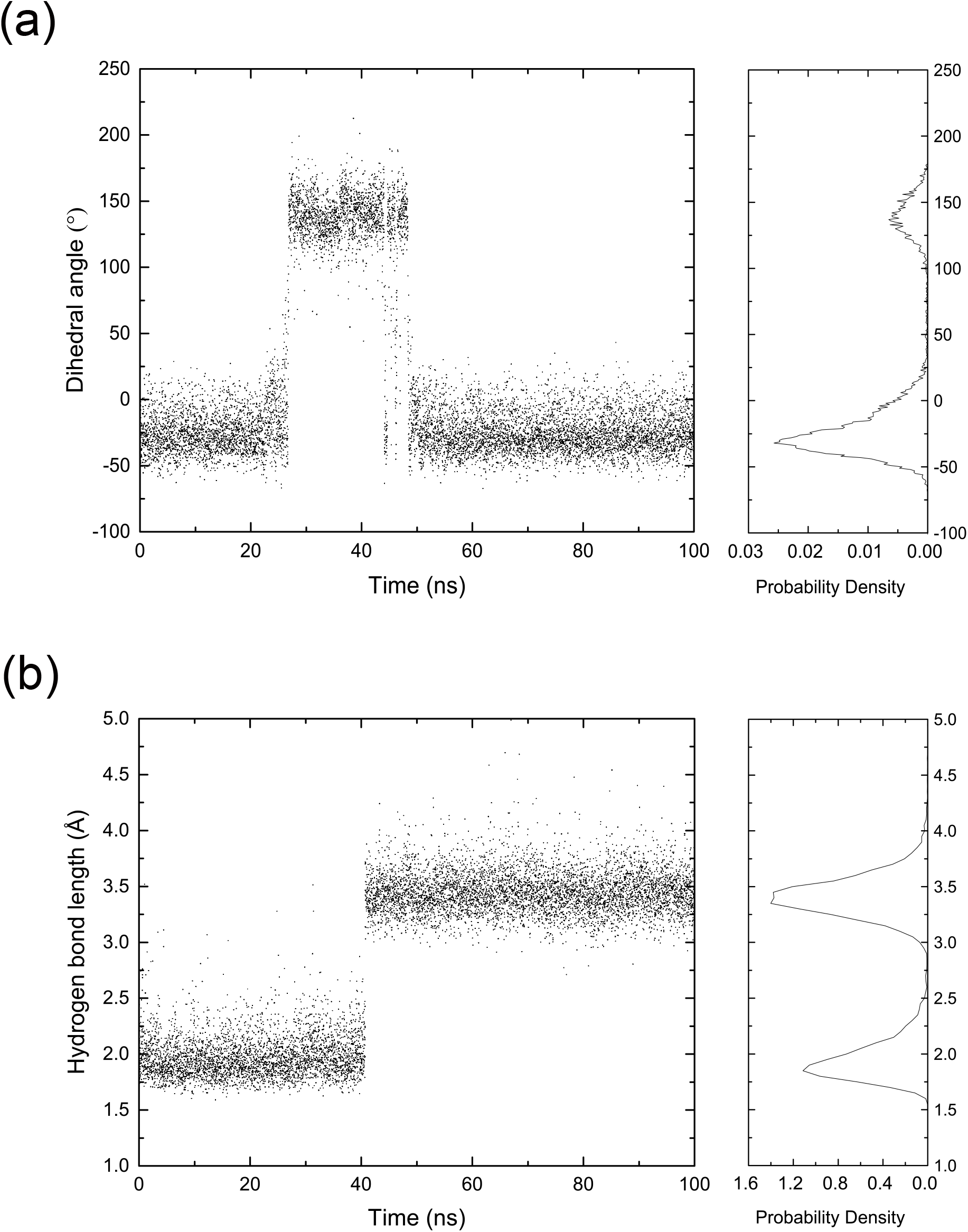
Examples of the behaviors of the boost targets during a transition and their probability density distributions; (a) dihedral angle and (b) hydrogen bond. Each variable fluctuates around clearly distinguishable values before and after the transition so that the resultant probability distribution has two peaks.

The forces acting on the four atoms *a, b, c*, and *d* by the bias potential *V*_*b,ϕ*_(*ε*_*ϕ*_) for the dihedral angle *ϕ* are

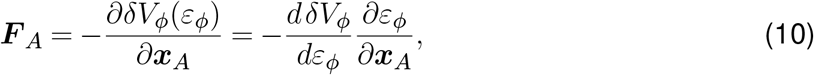

where *F* _*A*_ is the force acting on a representative atom *A* (=*a, b, c*, or *d*), *x*_*A*_ is the position vector of atom *A*, and

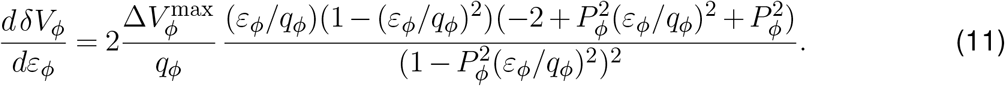

The dihedral angle *ϕ* is defined by the four atom positions as

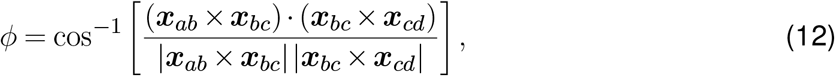

where *x*_*ab*_ = *x*_*b*_ *− x*_*a*_, *x*_*bc*_ = *x*_*c*_ *− x*_*b*_, and *x*_*cd*_ = *x*_*d*_ *− x*_*c*_. Thus, the forces applied to the four atoms are as follows:

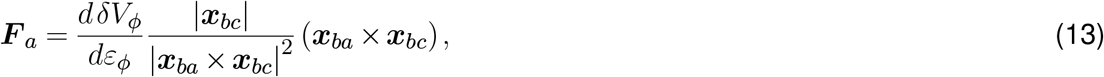

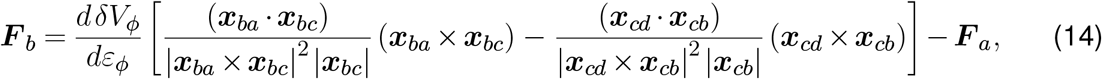

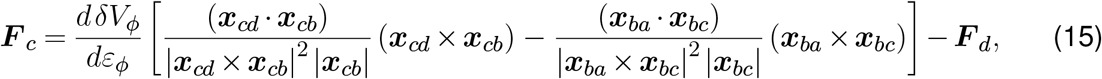

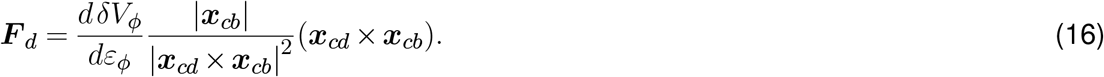

In a similar way, the forces acting on the two atoms *a* and *b* by the bias potential *δV*_*r*_(*ε*_*r*_) for the hydrogen bond *r* = |*x*_*a*_ *− x*_*b*_| are

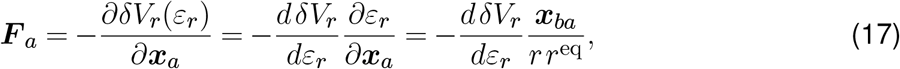

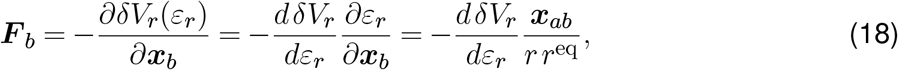

where *r*^eq^ is the equilibrium length of the hydrogen bond, and *dδV*_*r*_*/dε*_*r*_ is similar to Eq. (11).

### Algorithm

A standard algorithm of our modified BBM is presented in Figure 3. The algorithm can be divided into three processes:

- Preparation: The information about hinge regions and hydrogen bonds should be prepared before starting the simulation.
- Setup: Dihedral angles and hydrogen bonds that meet the conditions based on the prepared information are tagged as a boost list, and the bias potential is applied only to them during the simulation. This is reasonable because the tagged angles in hinge regions constituting a loop or linker and hydrogen bonds between the protein and the ligand are considered to be related to the transition. After all the boost bonds and angles are adopted, a conventional MD simulation is carried out for *t*^waiting^ to measure the average values of the boost list. The average value of each boost target is finally chosen as the equilibrium position.
- Implementation: The algorithm uses the threshold at the start of each step to determine whether the system is in the current state or has moved out of it. If a boost target is below its threshold value, a bias force is applied, and the equilibrium position is updated every 100 ps by averaging the values over this interval to compensate for changes in the internal local state. In contrast, if a boost target becomes greater than its threshold, its bias potential is immediately set to zero to protect the transition state. Lastly, if the boost target is out-of-state continuously and does not revisit the previous state for a certain period of time, it is assumed that a transition to a new state has occurred. In this case, an equilibrium position of the transited system is reestablished through the process described in Setup.

**Figure 3.**
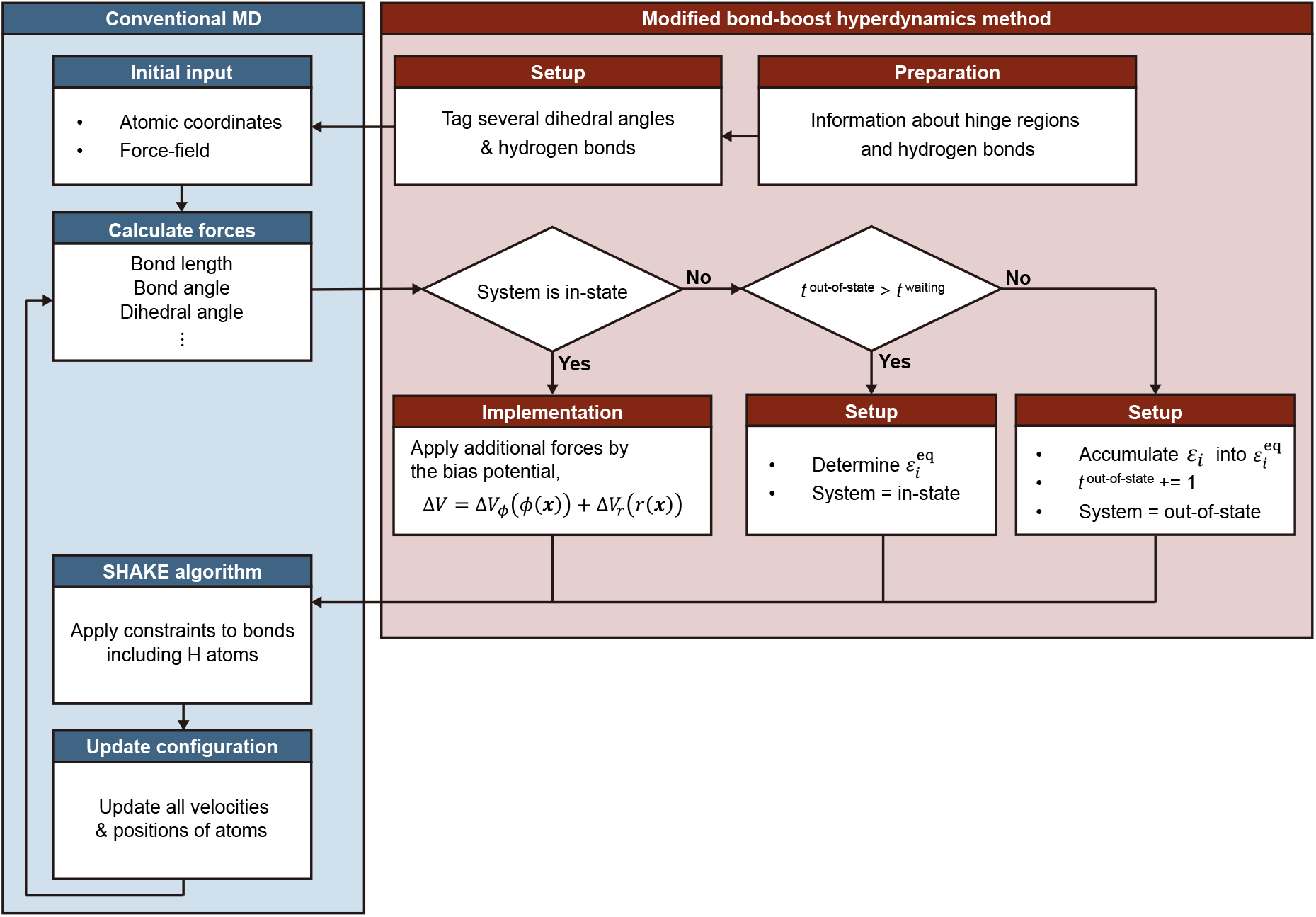
A flow chart of the algorithm of the modified bond-boost hyperdynamics method for investigating the conformational transition pathway of proteins. The bias force is computed, subsequent to the force calculation of the original force field, and the system’s state, whether in-state or out-of-state, is determined at each step using the predefined threshold. Our algorithm incurs a computational overhead of merely 10% of the total computational cost.

This algorithm was implemented as a package of the Large-scale Atomic/Molecular Massively Parallel Simulator (LAMMPS) program [35]. The added code runs after the calculation of the bonded and non-bonded interactions and before the SHAKE algorithm [36] in the LAMMPS simulation. Accordingly, it is ensured that the covalent bonds connected to H atoms are correctly constrained by the SHAKE algorithm when the bias force is added to the existing force. The computational overhead due to the added part for the modified BBM is only within 10% of the total computational cost of the MD simulation.

### Simulation details

To assess the reliability and validity of the developed method, we performed conventional and accelerated MD simulations for chain A of AdK’s closed structure (1AKE:A). The structure obtained from *Escherichia coli* was identified with X-ray diffraction and has P1, P5 bis(adenosine-5’-)pentaphosphate (Ap5A), which describes ATP+AMP, as an inhibitor [29]. For the system of 1AKE, Ap5A was removed, and two ADP molecules along with Mg^2+^ ion were obtained from *Mycobacterium tuberculosis* AdK structure (PDB ID: 2CDN) [37]. To integrate the ADP molecules and Mg^2+^ ion into the system, the adenosine and adjacent phosphate groups of the ADP were RMS-fitted to the corresponding atom positions of Ap5A [38].

For the closed structure with ADP · Mg^2+^ + ADP, the conventional and accelerated MD simulations were carried out three times using LAMMPS [35]. The AMBER ff14sb force field [39] was utilized for the protein, and the parameters of Meagher et al. [40] and Allnér et al. [41] were used for ADP and the magnesium ion, respectively. The system was solvated with the explicit TIP3P water model [42] in a cubic box of 75*×*75*×*75 Å^3^ and 8 Na^+^ ions were added to neutralize the system. All the covalent bonds involving hydrogen atoms were constrained by the SHAKE algorithm. An MD time step of 2 fs was used. All the simulations were integrated with the Verlet algorithm [43] and performed with the Nosé-Hoover thermostat and barostat [44, 45, 46]. The potential energy minimization of the system was performed using the conjugate gradient method, and three different systems were generated by varying the initial velocity. For equilibration, 200 ps NVT simulations at 300 K and 1000 ps NPT simulations at 300 K and 1 atm were carried out. Finally, 100 ns production runs of the NPT simulations at 300 K and 1 atm were conducted for both conventional and accelerated MDs on the three systems.

The simulation results were analyzed using the VMD program [30] to visualize and interpret the data. In particular, the distances of salt bridges were calculated using the salt bridge plugin of VMD, which measures the distance between a basic nitrogen and an acidic oxygen, and we judged that interactions were broken when they exceeded 6 Å. To evaluate the formation of salt bridges, we assessed the probability of formation at 0.2 Å intervals for both the CORE–LID and CORE–NMP distances. In addition, the umbrella sampling method was used to calculate the two-dimensional free energy landscape of AdK with the CORE–LID distance and the CORE–NMP distance as two reaction coordinates [5]. The two-dimensional domain ranges between 18–34 Å for the CORE–LID distance (the *x* dimension) and between 17–25 Å for the CORE–NMP distance (the *y* dimension), respectively. A total of 78 windows were created by splitting the *x* and *y* dimensions by 1.0 Å intervals. A harmonic restraint with a spring constant of 5 kcal/mol/Å^2^ was applied to each window and the accelerated MD trajectory was used as a starting structure of each window. However, note that the sampling result is independent of the initial configuration so that the resultant free energy surface is constructed without any bias. Each window simulation was performed for 5 ns (2,500,000 MD steps), and its trajectory was sampled every 10 steps except for the initial 500 ps. Therefore, 225,000 samples were collected for each window, and then the potential of mean force was calculated using the weighted histogram analysis method (WHAM) [47].

## Results and discussion

Conventional and accelerated MD simulations starting from the closed structure were performed three times for 100 ns each. During the simulation, a significant difference in the behavior of AdK was observed between them, confirming the performance of our approach. Furthermore, the conformational transition pathways of AdK observed in the accelerated MD simulations were compared with previously reported results to verify the reliability of our approach. In particular, distinct domain behaviors were identified in the release process of AdK across the three systems during this analysis, suggesting the involvement of multiple pathways. Interdomain salt bridges were also investigated at the atomic level to assess the stability of the generated structures (i.e. intermediate and open structures). Finally, umbrella sampling method and WHAM were applied to the trajectories of the three systems to determine the free energy landscape of AdK.

### Root mean square deviation (RMSD)

To monitor the conformational change of AdK, we measured the root mean square deviation (RMSD) with respect to the open structure 4AKE:A for C_*α*_ atoms (Figure 4). Since the systems start with the closed-structure, initially the RMAD values fluctuate around higher levels (6 to 8 Å) and is expected to decrease approaching zero if the protein transitions to the open structure. As clearly seen in the figures, the RMSD of the conventional MD simulations remain at higher values in all of the three systems, indicating the systems remain in their initial closed structure, which is consistent with previous studies [48, 49]. In contrast, the RMSD in the accelerated MD simulation of System 1 (Figure 4a) decreases to 4 Å within the first 15 ns and subsequently drops to about 2 Å by 40 ns. The accelerated MD simulations of Systems 2 and 3 also observed similar decrease in RMSD to sufficiently lower values within 50 ns (Figures 4b and 4c). These changes in RMSD observed in the accelerated MD simulations suggest that the proteins in the three systems all transitioned to the open structure.

**Figure 4.**
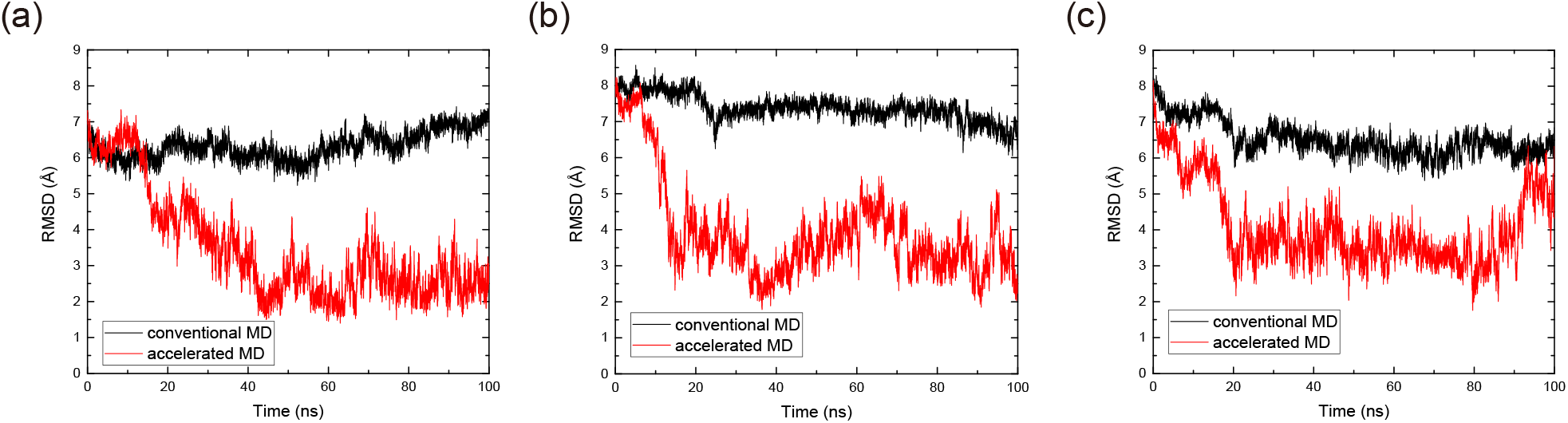
RMSD changes over time, comparing the conventional (black) and accelerated (red) MD simulation results for the C_*α*_ atoms in (a) System 1, (b) System 2, and (c) System 3. The RMSD was measured with respect to the open structure (4AKE), whereas the systems started with the closed structure.

Actual conformational changes of AdK were directly observed by monitoring the changes in atom configuration as shown in Figure 5. Figure 5 is an expanded version of Figure 4a, provided to offer a more detailed view of the conformational changes in System 1. In case of System 1, throughout the transition, the states change sequentially as each domain moves: initially, both the LID and NMP are closed; then, the LID opens while the NMP remains closed; and finally, the NMP opens. In fact, System 3 exhibited similar behavior to System 1, whereas System 2 followed a different sequence, with NMP opening first, followed by LID, leading to the transition to the open state. These observations suggest that during the closed-to-open transition, AdK can undergo multiple conformational pathways. These results clearly demonstrate that accelerated MD simulation, as implemented with our method, significantly outperforms conventional MD simulation in overcoming the conformational energy barriers. To further investigate these pathways and the domain motions in greater detail, additional analyses were conducted in the next section.

**Figure 5.**
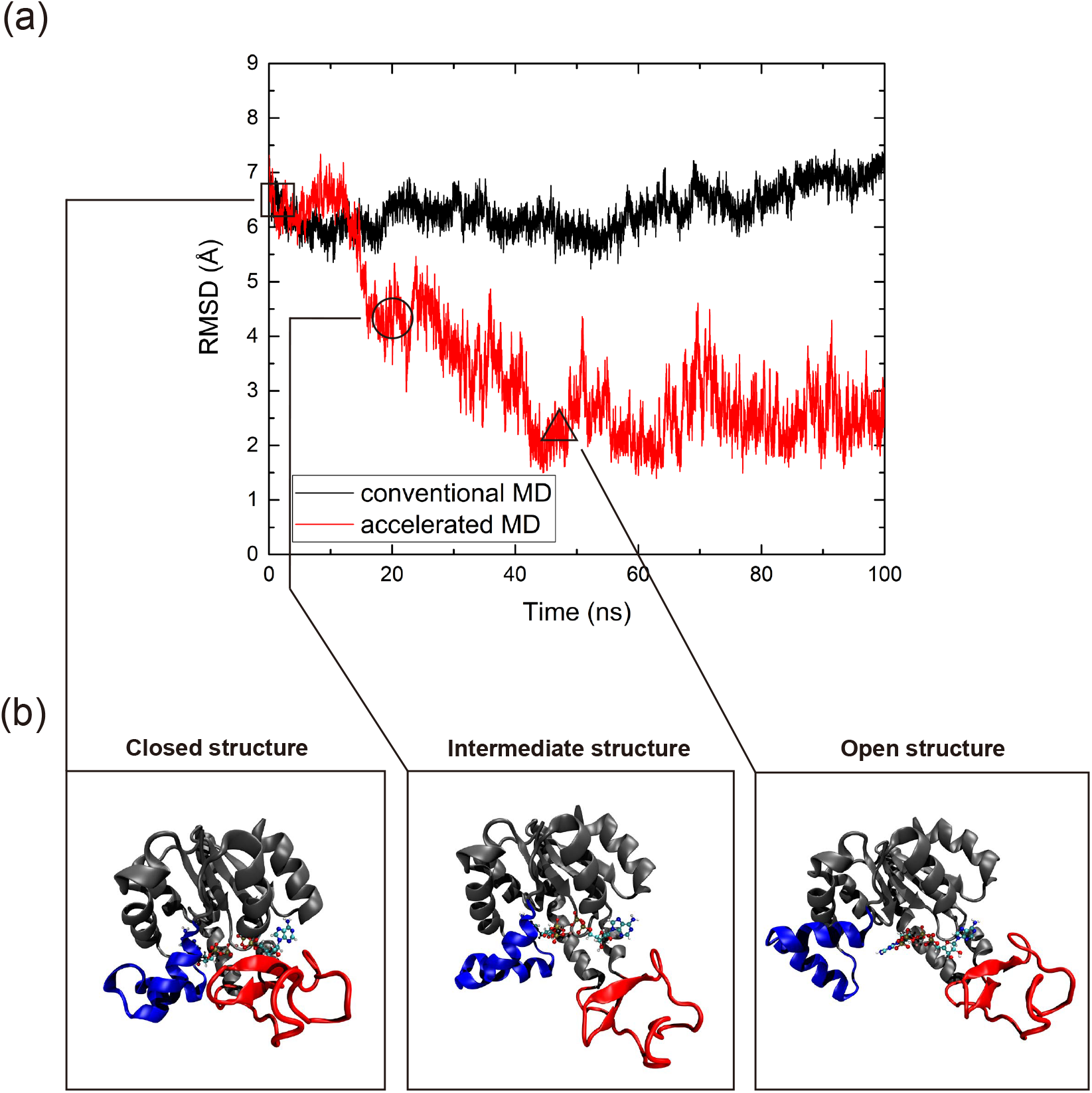
Comparison between the conventional and accelerated MD simulation results of System 1; (a) RMSD changes over time. The RMSD in the conventional MD remains fluctuating around 7 Å for 100 ns while the RMSD in the accelerated MD exhibits an initial drop to around 4 Å and finally decreases below 2 Å around 40 ns. (b) The conformational change of AdK in the accelerated MD over time. It starts with the closed structure (LID-closed/NMP-closed, symbol □), changes to the intermediate structure (LID-open/NMP-closed, symbol ○), and then reaches the open structure (LID-open/NMP-open, symbol Δ), sequentially.

### Conformational transition pathway

To elucidate the domain movement more specifically, we considered two other variables: the CORE–LID and CORE–NMP distances. The CORE–LID distance is defined as the distance between the center-of-masses of the LID and the CORE domains, and similarly, the CORE–NMP distance is defined as the distance between the center-of-masses of the NMP and the CORE domains. Because two ADP substrates bind between the CORE and the LID and between the CORE and the NMP, the conformational transition can be accurately described by these representative variables. These two variables are plotted as a two-dimensional map in Figure 6 with the CORE–LID and CORE–NMP distances designated as the *x* and *y* axes, respectively. In the map, the trajectories of the conventional and accelerated MDs are represented by black dots and red dots, respectively, and both simulations start from the lower left position. The two blue dots in the lower left corner and in the upper right corner also indicate the closed structure (1AKE:A) and the open structure (4AKE:A), respectively. As observed in Figure 6a, AdK of the accelerated MD undergoes a sharp increase in the CORE–LID direction, reaching to approximately 30 Å, followed by a subsequent rise in the CORE–NMP direction from 18 to 24 Å. System 3 also followed a similar pathway (Figure 6c), and these results indicate that, during the release process of AdK, the LID domain moves toward the open state first, followed by the NMP domain. However, as noted above, in System 2, the NMP domain opened first, resulting in a distinct pathway compared to System 1 (Figure 6b). This divergent behavior suggests the presence of multiple conformational pathways for the release process of AdK, in agreement with previously reported findings [49]. Specifically, based on the accelerated MD of System 1 (Figure 6a), it is expected that two transition states exist, dividing the system into three states. The first transition state is located near the position (25, 18.5) Å, and the second one is at the CORE–NMP distance of 19 Å between the CORE–LID distances of 27 and 30 Å. The states separated by these transition states represent the closed, intermediate (LID-open/NMP-closed), and open structures, identified in the above RMSD graph (Figure 5). This transition pathway is also consistent with those reported in previous studies [48, 50]. In contrast to the accelerated MD simulations, all the conventional MD simulations show that AdK remains trapped in the initial state and cannot escape during the simulation.

**Figure 6.**
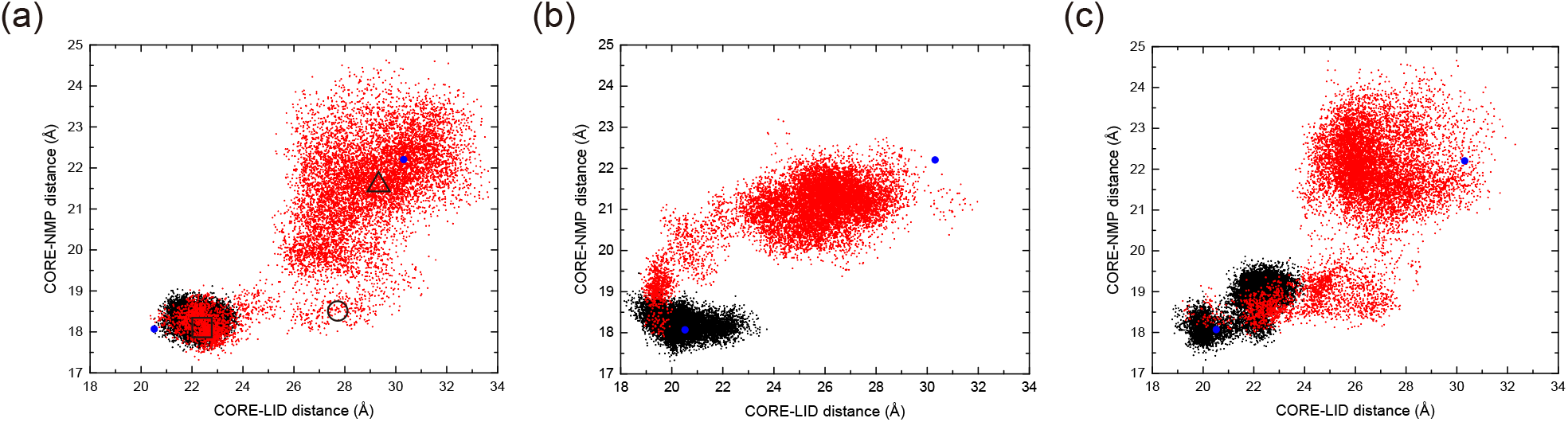
Two-dimensional projection of the CORE–LID and CORE–NMP distances for the conventional (black) and accelerated (red) MDs for (a) System 1, (b) System 2, and (c) System 3. The blue dots located in the lower left and upper right corners correspond to the closed (PDB ID: 1AKE) and open (PDB ID: 4AKE) structures of AdK, respectively. In panel (a), the □, ○, and △ symbols represent the closed, intermediate, and open structures shown in Figure 5, respectively. Both the conventional and accelerated MD simulations begin at the lower left corner.

For System 1, the two representative distances were also plotted against time in Figure 7. Similar to the above results, the plot shows a change in the CORE–LID distance from 22 to 30 Å within 15 ns (Figure 7a), followed by an increase in the CORE–NMP distance to approximately 23 Å at 40 ns (Figure 7b). In conclusion, these results demonstrate that the accelerated MD not only explores wider regions in an accelerated pace, but it also accurately follows the natural conformational transition pathway of AdK.

**Figure 7.**
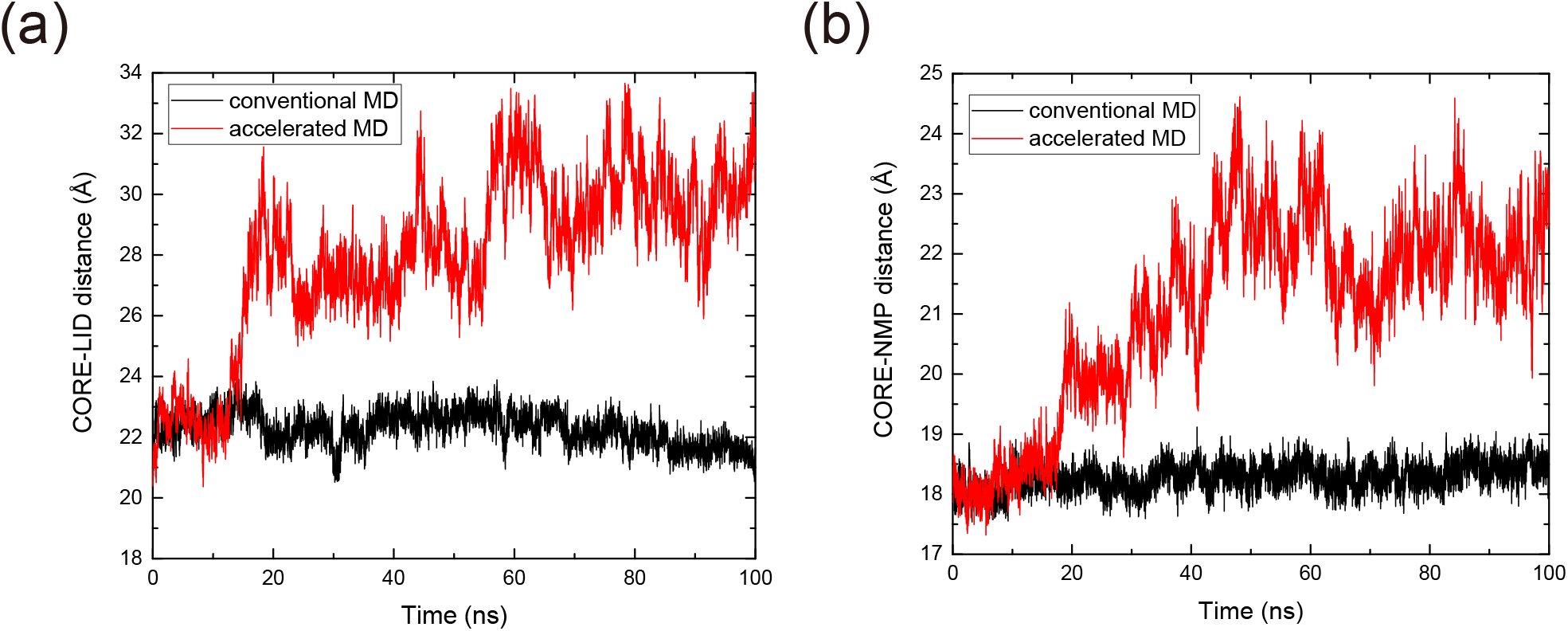
The time-dependent changes in distance between the domains in the conventional and accelerated MDs. (a) The distance between the center-of-masses of the LID and CORE domains. In the conventional MD, the distance remains relatively constant throughout the simulation, while in the accelerated MD, it changes before 20 ns and then fluctuates around 26–30 Å. (b) The distance between the center-of-masses of the NMP and CORE domains. In the accelerated MD, the distance increases significantly around 30 ns, after which AdK moves freely in the open state.

### Salt bridges

In addition to investigating the conformational transition pathway, we also conducted an analysis to evaluate the structural stability of AdK during the large conformational changes at the atomic level. In AdK, the structure with a number of charged residues is stabilized by various interactions including hydrogen bonds and salt bridges. Particularly, interdomain salt bridges are formed between different domains and undergo characteristic changes along with the transition [23]. To ensure that the structures have been properly generated, we monitored the formation of three interdomain salt bridges in AdK: K57–E170 (CORE–NMP), R36–D158 (LID–NMP), and D118–K136 (CORE–LID).

Using the accelerated MD result of System 1, we investigated the behavior of three salt bridges in response to conformational changes of AdK, as depicted in Figure 8. Our results indicate that salt bridges that are typically conserved in the closed structure, such as K57–E170 and R36–D158, are disrupted during the transition. Specifically, Figures 8a and 8b show the interaction formation probability of the three salt bridges as a function of the CORE–LID and CORE–NMP distances, respectively. In Figure 8a, the K57–E170 and R36–D158 salt bridges exhibit a similar trend of disruption with increasing the CORE–LID distance, but it is clear that R36–D158 is broken first, followed by K57–E170. A similar trend is observed in Figure 8b, where the salt bridge between LID-NMP, R36–D158, is broken first, reflecting the aforementioned transition pathway in which the LID domain opens first. In contrast, the D118–K136 salt bridge behaves differently from K57–E170 and R36–D158; it forms an interaction as the structure transitions to the open state (Figures 8a and 8b). Given that the D118–K136 interaction plays a critical role in stabilizing the open structure [25], this result suggests that a stable open state is achieved once the LID domain is sufficiently open.

**Figure 8.**
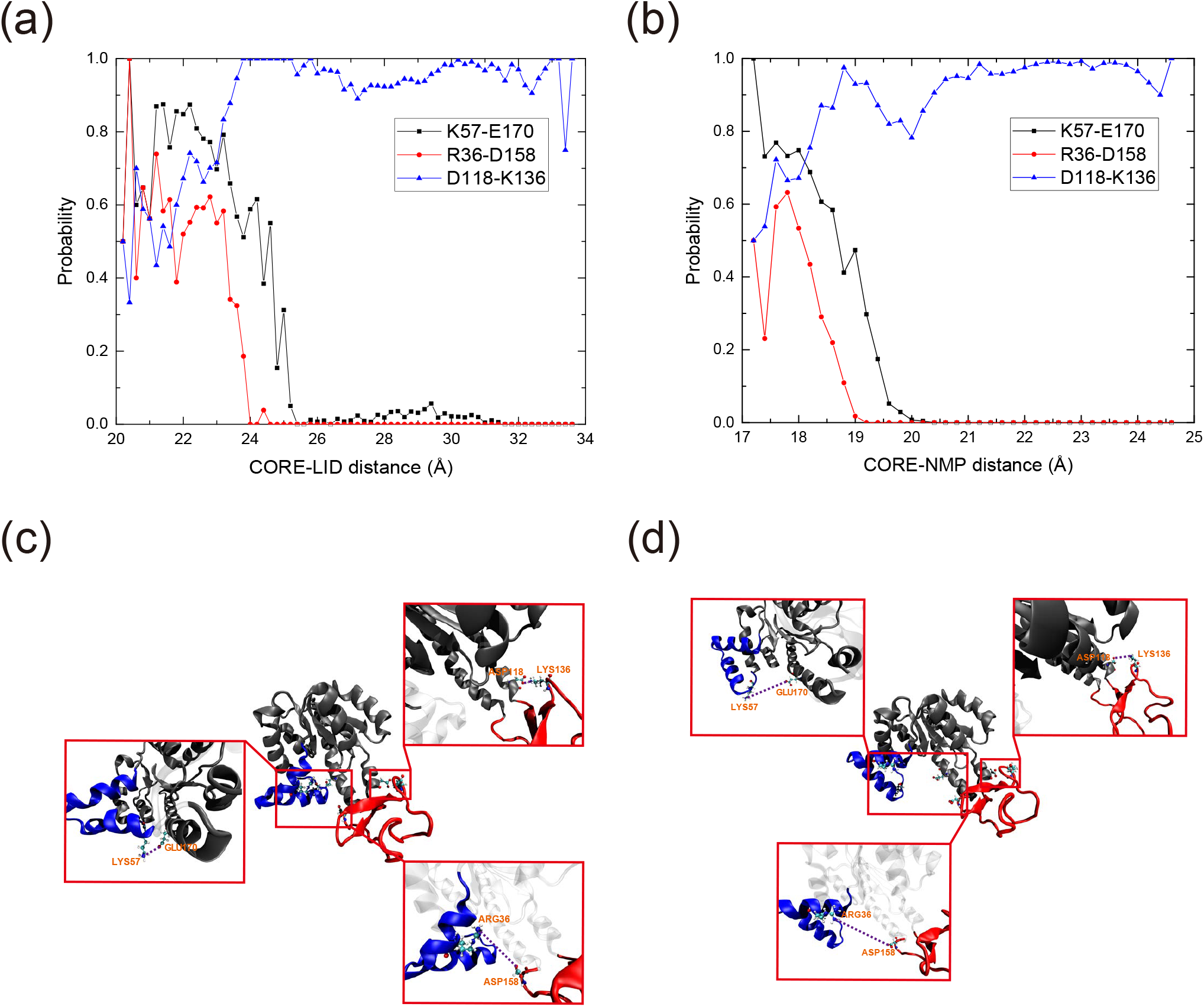
Probability changes for salt bridge formation in two collective variables. The probability was calculated by dividing each collective variable at an interval of 0.2 Å. (a) Probabilities of K57–E170, R36–D158, and D118–K136 formations according to the CORE–LID distance. (b) Probabilities of K57–E170, R36–D158, and D118–K136 formations according to the CORE–NMP distance. (c) A snapshot of the three salt bridges in the intermediate structure. (d) A snapshot of the three salt bridges in the open structure.

This is further confirmed in the intermediate structure (Figure 8c) and the open structure (Figure 8d). In Figure 8c, the R36–D158 salt bridge is disrupted as the red LID domain transitions to the open state, while the blue NMP and gray CORE domains remain closed, allowing the K57–E170 interaction to persist in the intermediate state. The K57–E170 salt bridge is subsequently disrupted once the structure fully transitions to the open state (Figure 8d). Meanwhile, as shown in Figures 8c and 8d, the D118–K136 salt bridge remains intact as the LID domain opens from the intermediate to the open structure. Through our investigation of the salt bridge formation/disruption processes, we have confirmed that the mechanism of AdK previously described in the literature is consistent with our findings. This further supports that the target protein behaves appropriately at the atomic level during accelerated MD simulations.

### Free energy landscape

The free energy landscape was calculated as a function of the two reaction coordinates, the CORE–LID distance and the CORE–NMP distance, by applying umbrella sampling combined with WHAM [47]. The entire domain was divided into 105 subdomains by placing 15 intervals along the CORE–LID direction and 7 intervals along the CORE–NMP direction. A total of 78 window simulations were performed, each exclusively sampling a single subdomain. Also, note that some subdomains were not explored because they are close to regions with relatively high free energies so that the system is unlikely to visit them. The result is shown in Figure 9. First, we found that a free energy minimum is located near the coordinate corresponding to the closed structure (the starting configuration) at CORE–LID = 22.5 Å and CORE–NMP = 18.5 Å. This finding together with the distribution of the simulation points in the CORE–LID distance and CORE–NMP distance domain shown in Figure 6 indicates that the conventional MD simulation was unable to overcome the energy barriers surrounding the closed state, so it remained around its starting configuration. In contrast, the accelerated MD simulations successfully crossed these energy barriers, transitioning to the adjacent intermediate states. The intermediate structures spanned a wide range of conformational space, reflecting the existence of multiple pathways. Notably, Systems 1 and 3 transitioned to the open structure via the downward path, while System 2 followed the upward path, consistent with the transition pathway results discussed above. After transitioning to the intermediate state, Systems 1 and 3 crossed the energy barrier in the CORE–NMP direction from the intermediate state, while System 2 moved in the CORE–LID direction. Eventually, all three systems reached the open structure, which has the lowest free energy value among the three structures. In the open state, the LID and NMP domains exhibited significant flexibility, which is attributed to the elimination of structural constraints imposed by interactions between ADPs and the domains, allowing for a wide range of conformational variability. In summary, the closed, intermediate, and open states visited by the accelerated MD simulations indeed correspond to the free energy minima, and our accelerated MD method effectively preserved the natural transition pathway in the free energy landscape of the original system, enabling the exploration of a wider conformational space.

**Figure 9.**
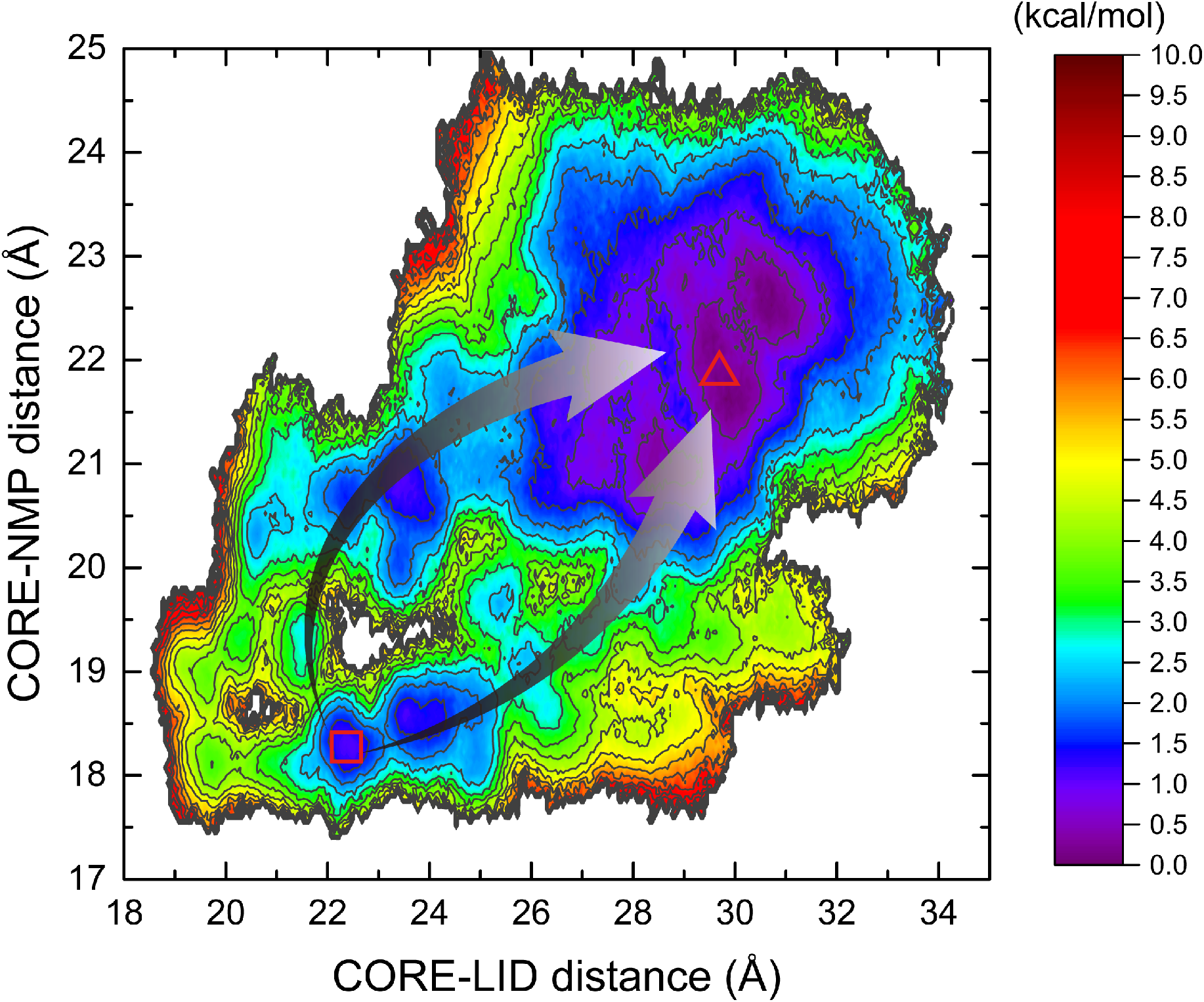
Free energy landscape of AdK constructed using the umbrella sampling method with respect to the CORE–LID distance and the CORE–NMP distance. The free energy at the minimum of the open structure was set to 0 kcal/mol and the contour lines are displayed at 0.5 kcal/mol intervals. Two pathways from the closed (symbol □) to the open (symbol △) structure are indicated by arrows.

## Conclusion

Biomolecular systems spend most time in their local minima before making transitions to other minima and thus biological phenomena are observed as rare events, making it difficult to be handled in MD simulations with the limited time-length. To address this issue, we employed an accelerated MD method with novel modifications of the BBM. Our bias potential was designed with new boost parameters such as the dihedral angle and the hydrogen bond to properly describe the conformational change of proteins and also removed the force discontinuity issue at the boosted and non-boosted boundary region. Since the bias potential was applied only to the basin excluding the transition state of the system, the natural conformational transition pathway was preserved while expediting the escape from the basin. In addition, an efficient algorithm to the method was developed and implemented as a LAMMPS package.

We validated the reliability of the developed method by applying it to the AdK system. First, we observed that the RMSD changed rapidly in the accelerated MD simulations, indicating the transition from the closed to the open structures of AdK. However, no such changes were found during the conventional MD simulations. Secondly, the closed-to-open transition was tracked as the movements of three domains, i.e. the LID, NMP, and CORE domains in AdK structure. In the accelerated MD simulations, two pathways were identified according to the sequence of domain opening, ultimately leading to the fully open structure of AdK. These multiple pathways have also been reported in previous studies. In addition, we observed that the salt bridges R36–D158 and K57–E170 were broken during the closed-to-open transition, while the salt bridge D118–K136 was formed in the open structure. By monitoring the changes in the salt bridge interactions, we confirmed that our technique accurately represented the atomic level movement of the protein. Finally, the free energy landscape of AdK was determined by applying umbrella sampling with WHAM. The closed, intermediate, and open states found in the accelerated MD trajectory were all expressed as local minima in the free energy landscape, and the energy barriers formed between them were also confirmed. These positive results prove that our accelerated MD technique will shed light on the time scale problem of MD simulations with broader applications in various biomolecular systems.

## Acknowledgements

This work was supported by Basic Science Research Program through the National Research Foundation of Korea (NRF) funded by the Ministry of Education (2021R1A6A1A03039696).

